# Varicella-zoster virus CNS vasculitis and RNA polymerase III gene mutation in identical twins

**DOI:** 10.1101/244848

**Authors:** Madalina E Carter-Timofte, Anders F Hansen, Maibritt Mardahl, Sébastien Fribourg, Franck Rapaport, Shen-Ying Zhang, Jean-Laurent Casanova, Søren R Paludan, Mette Christiansen, Carsten S Larsen, Trine H Mogensen

## Abstract

Deficiency in the cytosolic DNA sensor RNA Polymerase III was recently described in children with severe varicella zoster infection in the CNS or lungs. Here we describe a pair of monozygotic female twins, who both experienced severe recurrent CNS vasculitis caused by VZV reactivation. The clinical presentation and findings included recurrent episodes of headache, dizziness, and neurological deficits, cerebrospinal fluid with pleocytosis and intrathecal VZV antibody production, and magnetic resonance scan of the brain showing ischaemic lesions. We performed whole exome sequencing and identified a rare mutation in the Pol III subunit *POLR3F*. The identified R50W *POLR3F* mutation is predicted to be damaging by bioinformatics and when tested in functional assays, patient PBMCs exhibited impaired antiviral and inflammatory responses to the PoL III agonist Poly(dA:dT) as well as increased viral replication in patient cells compared to controls. Altogether, these cases add genetic and immunological evidence to the novel association between defects in sensing of AT-rich DNA present in the VZV genome and increased susceptibility to severe manifestations of VZV infection in the CNS in humans.

**Abbreviations:**

CADD: combined annotation dependent depletion
CSF: cerebrospinal fluid
DOCK: dedicator of cytokinesis
IFNGR: interferon gamma receptor
MSC: mutation significance cut-off
NK: natural killer
POLIII: RNA polymerase III
SCID: severe combined immunodeficiency
TYK: tyrosine kinase
VZV: varicella zoster virus
WES: whole exome sequencing

## Introduction

Varicella zoster virus (VZV) is a human pathogenic alpha-herpesvirus causing chickenpox in children during primary infection and herpes zoster in elderly or immunocompromised individuals upon reactivation from latency. However, in a minority of infected individuals, VZV may cause pneumonia or even more rarely infection in the central nervous system (CNS). VZV disease manifestations in the CNS may present as a viral meningoencephalitis with classical signs of viral meningitis but may also present in a more atypical stroke-like manner, in which case the immunopathogenesis appears to be vasculitis ^(1)^. VZV vasculitis can occur during primary infection or during reactivation from latency, and in both cases with considerable delay, and may affect either small-or large vessels of the cerebral parenchyma. Whereas large-vessel disease is most common in immunocompetent individuals, small-vessel disease usually develops in immunocompromised patients; however, in some patients both large and small vessels are involved. Overall, CNS involvement in VZV infection, of which CNS vasculitis only represents a smaller fraction, is very rare with an estimated incidence of 1-3/10.000 primary VZV infections, whereas the incidence of meningoencephalitis during VZV re-activation is more difficult to establish ^(2;3)^.

Here we describe two monozygotic twins with similar clinical presentations suggesting recurrent CNS vasculitis caused by VZV reactivation. Genetic and functional immunological analyses were performed in order to examine a possible genetic and pathophysiological basis of the clinical phenotype.

## Methods

### Patient material

The patient and her twin sister were admitted for clinical immunological evaluation. See supplementary medical history for further clinical details. A total of 84 ml of blood was drawn from each patient. 4 ml were used for DNA-isolation and subsequent WES, whereas the remaining 80 ml were used for isolation of peripheral blood mononuclear cell (PBMC)s, performed by using Sepmate tubes (Stemcell Technologies) with Ficoll-plaque (GE Healthcare Lifesciences) and cells were then stored in liquid nitrogen. Control PBMCs were obtained from healthy controls after written consent.

### Whole Exome Sequencing (WES)

DNA was isolated from EDTA stabilized blood using EZ1 DNA Blood 350 µl Kit and an EZ1 Advanced XL instrument (Qiagen) according to manufacturer’s instructions. See supplementary methods for description of WES data analysis.

### Stimulation of PBMCs

PBMC’s from patients and controls were thawed in 50 ml tubes containing 15 ml of preheated media (RPMI-1640 w/L-glutamin (Biowest)) supplemented with 10% heat-inactivated fetal bovine serum (Biowest) and 1% penicillin/streptomycin and spun down at 350g for 8min. The PBMC’s were resuspended in media and divided into 24-well plates at a concentration of 5 × 10^5^ cells per 300 μl media per well. Next, cells were incubated overnight at 37 °C in an atmosphere of 5% CO_2_. The cells were subsequently transfected using Lipofectamine3000 at 0,75μl/μg DNA (Invitrogen by Thermo Fischer Scientific) and Opti-MEM (Gibco, by Life Technologies by the TLR3-agonist Poly(I:C) (2 μg/mL) or transfected with the RNA-polymerase III agonist Poly(dA:dT) (2μg/mL) (Cayla, Invivogen). Cells were incubated for 6 hours before harvest and cell lysis. For virus experiments PBMCs were infected with VZV-infected MeWo cells (ROka strain) (PBMC:MeWo-VZV ratio, 1:1). One vial of VZV-infected MeWo cells was thawed, spun down at 1000 rpm for 10 min and re-suspended in 400 µL preheated media. From this solution, 30 µL was added to the wells to be stimulated with VZV. Cells were incubated for 6 h, except VZV which were incubated with PBMCs for 48 h before cells lysis and RNA harvest. The stimulations were performed in triplicates and each experiment was performed 2–3 times.

### Isolation of RNA and RT-qPCR

RNA was purified from PBMC whole-cell lysates, as per manufacturer instructions, using High Pure RNA Isolation Kit (Roche). Prior to cDNA synthesis, VZV-infected PBMCs underwent DNAse treatment and removal step (Ambion, Thermo Fisher Scientific). From the isolated RNA, cDNA was synthesized by using QuantiTect Reverse Transcription Kit (Qiagen) following the manufacturer’s instructions. The synthesized cDNA was subsequently used for real time qPCR using TaqMan probes, allowing amplification and analysis of levels of *IFNB1*, *TNFA*, the interferon (IFN)-stimulated gene *CXCL10*, and the viral VZV gene *ORF63*. *TBP* was used as housekeeper gene for reference. All analyses were performed as technical duplicates for all samples and the TaqMan probes (Life Technologies) used were: *IFNB1*: Hs01077958, TNFA: Hs01113624, *CXCL10*: Hs01124251. For analysis of *ORF63*, which was performed separately, SYBR Green was used for *ORF63* (LGC Biosearch Technologies).

### Statistical analysis

The Mann-Whitney rank t-test was used to determine statistical significance; non significant (ns); * p ≤ 0.05; ** p ≤ 0.001.

### Ethics

The project was approved by the Regional Ethics Committee (#1-10-72-275-15) and the patients provided informed written consent before blood sampling.

## Results

A 37 year-old female (P1) was referred to the international center for immunodeficiency diseases (ICID) for clinical immunological evaluation based on recurrent stroke-like manifestations diagnosed in the neurological department as CNS vasculitis caused by VZV reactivation. This diagnosis was based on clinical symptoms including headache, dizziness and hemiparesis, as well as paraclinical test revealing cerebrospinal fluid (CSF) pleocytosis, positive VZV IgG, and finally MR scan of the brain with ischaemic lesions. The monozygotic twin sister (P2) experienced a very similar medical history. For a detailed medical history and summary of paraclinical analyses, see Supplementary medical history and Table 1.

**Table 1.** Demographic-, genetic-, cerebrospinal fluid-, and clinical data for P1 and P2 with CNS vasculitis

| Patient | Age*, Gender | Gene | Transcript ID, transcript variant, protein variant | CADDscore (MSC) ExAC frequency | CSF-findings | MR-C findings | Clinical presentation Number of episodes | Treatment 1 Prophylaxis2 |
| --- | --- | --- | --- | --- | --- | --- | --- | --- |
| P1 | 25, female | POLR3F | NM_001282526.1<br>c.148C>T , p.R50W | 25.8 (13.0)<br>0.0000 | Pleocytosis<br>VZV PCR negative<br>VZV IgG positive | Repeated<br>ischemic<br>lesions/infarcts | Hemiparesis x 2<br>Paresthesias<br>Headache<br>4 episodes | 1) Aciclovir<br>+corticosteroid<br>2) Aciclovir |
| P2 | 35, female | POLR3F | As above | As above | Pleocytosis<br>VZV PCR negative<br>VZV IgG positive | Repeated<br>ischaemic<br>lesions/infacts | Paresthesias<br>Hemiparesis x 1<br>>4 episodes | 1) Aciclocir<br>2 ) Aciclovir |

\*Age at first clinical episode with neurological symptoms
CADD combined annotation dependent depletion; MSC mutation significance cutoff; CSF cerebrospinal fluid; MR-C magnetic resonance of cerebrum.

Due to this unusual presentation in both monozygotic twin sisters, we suspected a genetic etiology of the recurrent VZV CNS vasculitis. A routine clinical immunological evaluation was normal, including normal immunoglobulins, lymphocyte distribution and –proliferation within normal range, and a negative HIV test. We therefore performed whole exome sequencing (WES) on DNA from P1, which resulted in the identification of a novel missense mutation in the POLR3F subunit of the cytosolic DNA sensor RNA Polymerase III (Figure 1A). The mutation, R50W *POLR3F*, is rare and predicted to be damaging by bioinformatics software, such as Polyphen2 and SIFT and with a high combined annotation dependent depletion (CADD) score of 25.8, which is well above the mutation significance cut-off score (MCS) of 13.0 for this gene ^(4;5)^. Further population genetic analyses, based on study of *POLR3F* variations from the GnomAD database, demonstrate a very low frequency of non-synonymous and loss-of-function mutations with a CADD score >12, and a global minor allele frequency (MAF) < 10^−4^ for nonsense mutations. Moreover, when performing natural selection assessment based on mouse and human gene homologous comparison, the *POLR3F* gene appears to be under purifying selection as reflected by a dN/dS score of 0.0172 below the threshold of 1). Finally, there is a selective pressure neutrality index of 0.1264, predicting moderate purifying selection (Figure 1A and supplementary Figure 1).

**Figure 1.**
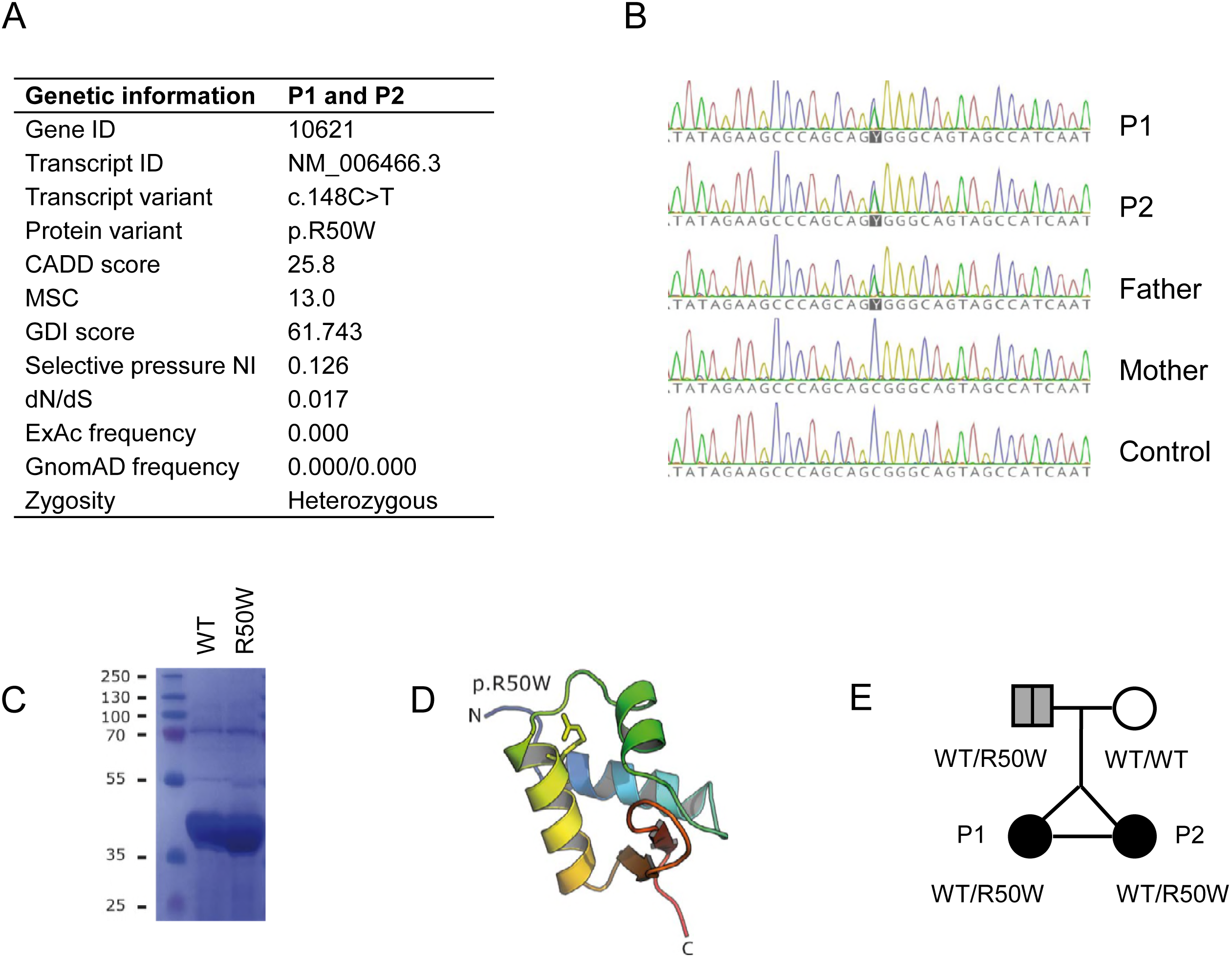
Identification of a heterozygous mutation in *POLR3F*, protein structure, and pedigree. A, Summary of genetic information on the identified variant in *POLR3F*. B, Sanger sequencing of P1, P2, their parents, and a control with the *POLR3F* variant c148C>T present in P1, P2 and the father. C, Coomassie-blue stain of SDS-PAGE revealing similar solubility of wild-type and mutant R50W *POLR3F* expressed from E.Coli. E, Molecular model of POLR3F first winged helix domain based on the Protein Databank ID 2DK8 with the R50 shown as sticks. D, Pedigree showing the affected monozygotic twins with the R50W *POLR3F* variant inherited in a heterozygous manner from the father with possible previous VZV CNS disease. CADD, combined annotation dependent depletion; MSC, mutation significance cut-of; GDI, gene damage index; NI, neutrality index.

The identified *POLR3F* variant (NM_001282526.1 c.25C>T, p.R50W) causes an amino acid shift from the positively charged arginine (R) to a bulky aromatic tryptophan (W) in the N-terminal region of the POLR3F subunit, which together with POLR3C and POLR3G constitutes a POLR3 sub-complex essential for POLR3 promoter transcription initiation(Figure 1A and D). As expected, the same POLR3F mutation was found by Sanger sequencing to be present in the monozygotic twin sister. The mutation was inherited from the father (Figure 1B), who was previously admitted to hospital with clinical signs and symptoms of stroke, which retrospectively might have been linked to VZV CNS vasculitis; however this was not examined at the time. Biochemical analyses demonstrated that expression and solubility of the mutant POLR3F protein was unaltered compared to wild-type, suggesting that the R50W mutation affects protein function, such as ligand recognition or enzymatic conversion of the AT-rich DNA ligand into 5’ triphosphorylated RNA, rather than expression of the POLR3F protein (Figure 1C). Two additional rare gene variants with a high CADD score identified through the WES analysis are listed in Supplementary table 2; however none of these were judged to be of any relevance for the infectious phenotype of these patients.

Next, we investigated in vitro antiviral and inflammatory responses in P1 to VZV infection and the Pol III agonist Poly(dA:dT), the latter being a constituent of the AT-rich VZV genome (Figure 2). As shown, PBMCs from P1 showed significantly impaired interferon (IFN)β –, TNFα –, and CXCL10 responses to poly(dA:dT) (Figure 2A-C). In contrast, responses to transfected Poly(I:C) were normal as expected, since this ligand activates the cytosolic RNA receptor RIG-I directly downstream of Pol III (Figure 2D-F). Moreover, we observed significantly impaired production of the two pro-inflammatory cytokines TNFα and IL-6 in response to VZV infection of PBMCs from P1 and controls (Figure 2J and K), although IFNβ, IFNλ, and CXCL10 responses were comparable between P1 and controls (Figure 2G-I). Importantly, upon VZV infection a significantly increased amount of the viral immediate-early gene *ORF63* mRNA was detected in P1 cells relative to controls, demonstrating increased viral replication, i.e. reduced viral control in patient cells (Figure 2L). Taken together, these results demonstrate impaired immune responses to the Pol III ligand Poly(dA:dT) and impaired pro-inflammatory responses to VZV in the setting of increased VZV replication in patient cells.

**Figure 2.**
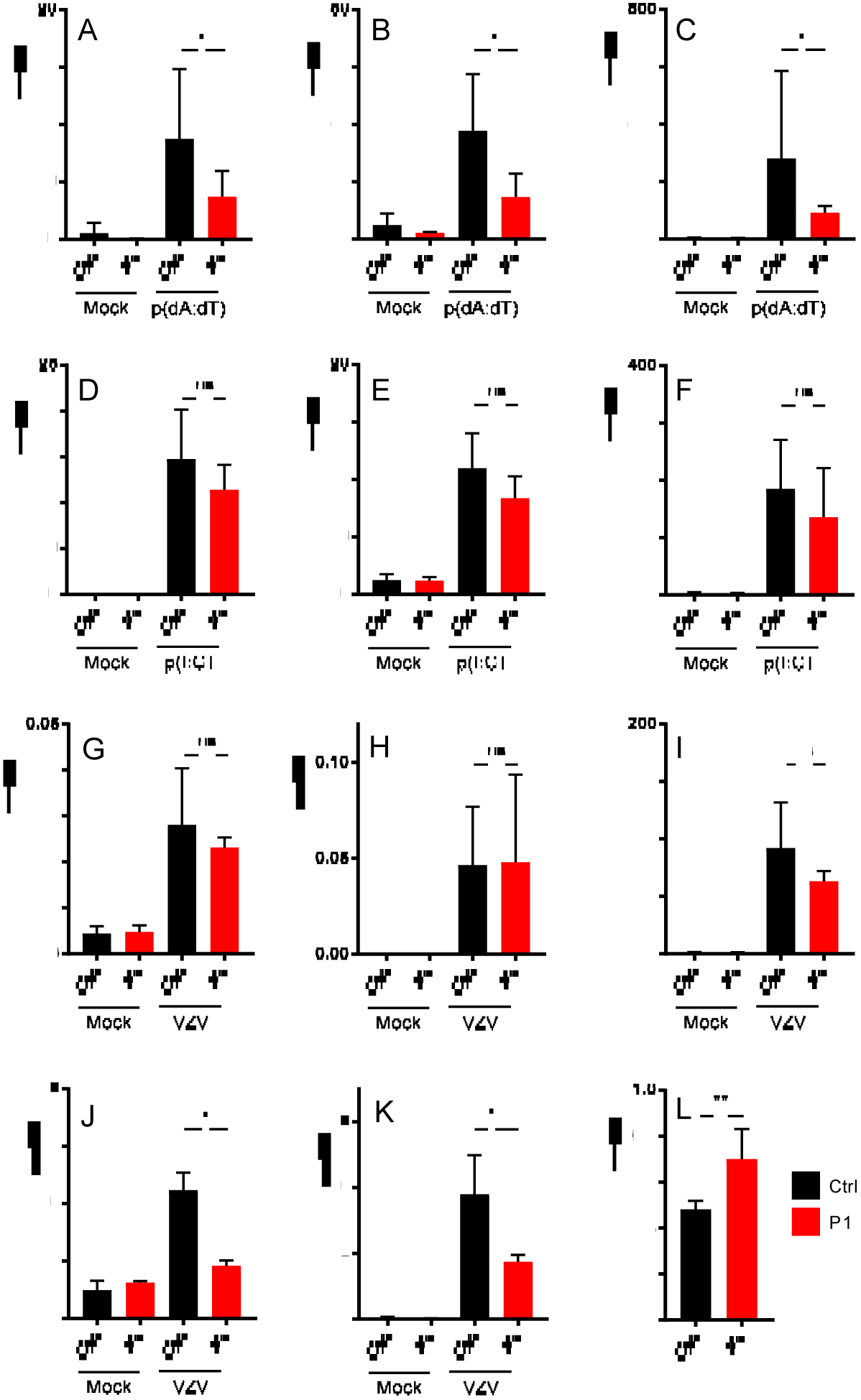
Impaired antiviral and inflammatory responses to the POL III ligand Poly(dA:dT) and increased VZV replication in patient PBMCs. PBMCs from P1 and nine healthy controls were used for experiments. A-C, PBMCs were transfected with poly(dA:dT) (2 μg/mL) for 6hs. D-F, PBMCs were transfected with poly(I:C) (2 μg/mL) for 6hs. G-L, VZV-infected MeWo cells were co-cultured with PBMCs for 48hs. Total RNA was isolated for measurement of mRNA for the indicated cytokines by RT-qPCR. L, VZV replication was accessed by measuring the viral (IE) transcript ORF63. Cytokine levels and ORF63 levels were normalized to TBP and compared to pooled results of nine healthy controls. Data are shown as column bars and error bars representing standard deviation. The non-parametric Mann-Whitney ranked sum test was used to evaluate statistical significance between groups. Significance level * p < 0.05 and ** p < 0.001.

## Discussion

A number of primary immunodeficiencies affecting innate and adaptive immunity have been previously recognized to predispose to severe disseminated VZV infection. These classically comprise disseminated VZV infection occurring in children with severe combined immunodeficiency (SCID), as well as immunodeficiencies more specifically involving defects in natural killer (NK) cells, including MonoMAC caused by *GATA2* mutation, but also involves a number of immunodeficiencies, in which the precise cellular and immunological basis of increased susceptibility to VZV infection is less well defined. These include autosomal recessive hyper-IgE syndrome caused by mutation in *DOCK8* or *TYK2*, Mendelian susceptibility to mycobacterial disease due to IFNGR1 deficiency, and finally, DOCK2 deficiency ^(6)^.

Recently, a novel primary immunodeficiency involving Pol III deficiency in four children with severe VZV infection in the CNS and/or lungs was described ^(7)^. In the present work, we provide further genetic and immunological evidence for a role of POL III in protective immunity to VZV infection, particularly in the CNS. RNA Polymerase III is a 17-subunit enzyme with dual functions in both promoter dependent transcription of tRNA and rRNA as well as in innate sensing of AT-rich DNA, converting this into 5’-phosphorylated single-stranded RNA, which serves as a ligand for the RNA sensor RIG-I ^(8;9)^. The present report is the first description of an association between the POLR3F subunit of Pol III and severe recurrent VZV CNS vasculitis. There has been one report of the presence of a potentially disease-causing mutant in *TLR3* in a 31-year old male patient with VZV meningoencephalitis, although no functional studies to support a role for TLR3 deficiency in VZV CNS infection were provided ^(10)^. Of note, mutations in the POLR3A and POLR3B subunits have been genetically linked to non-infectious hypomyelinating leukodystrophy ^(11;12)^.

The cases described here are notable for the occurrence of an extremely rare disease, ie CNS vasculitis caused by recurrent VZV reactivation in monozygotic twins. Although the presence of vasculitis could only be determined with certainty on MR angiography of the brain in P2, the clinical history consisting of recurrent neurological deficits presenting in a stroke-like manner with headache, hemiparesis, pleocytosis and elevated VZV IgG in CSF, white matter lesions detected on brain MR scan are all highly suggestive of this diagnosis. Indeed, in a case series of 14 adults with cerebral vasculitis, it was reported that positive intrathecal anti-VZV IgG combined with relevant neurological symptoms and imaging was a definite marker of this disease rather than VZV DNA detection by PCR, since the latter was positive in only four of the patients ^(13)^. In this respect, the absence of detectable VZV by PCR in CSF is not surprising and is in consistency with VZV CNS infection, including vasculitis ^(2)^. Importantly, both patients described in the present study have a good clinical response to acyclovir and corticosteroid treatment during clinical episodes but experience recurrence of symptoms and MR abnormalities during discontinuation of acyclovir. There is clearly a different pathogenesis between classical VZV meningoencephalitis and the presence of vasculitis developing in some cases of VZV CNS infection ^(1)^. At the cellular level, increased viral replication, which for VZV is known to also include endothelial cells within the CNS, may evoke enhanced inflammatory responses secondary to the presence of virus and thereby explain the occurrence of vasculitis and immunopathology in these patients.

## Conclusions

In summary, we describe two female monozygotic twins both experiencing recurrent stroke-like episodes diagnosed as CNS vasculitis caused by VZV reactivation, in whom we identified a heterozygous missense mutation in the POLR3F subunit of the cytosolic DNA sensor Pol III, resulting in impaired antiviral responses and increased VZV replication in patient PBMCs compared to controls. We propose that impaired sensing of AT-rich DNA in the VZV genome contributes to increased susceptibility to the rare condition of VZV-induced CNS vasculitis, thereby extending the spectrum of CNS manifestations linked to POL III deficiency. Altogether, this case adds further genetic and immunological evidence to the novel association between defects in DNA sensing and increased susceptibility to severe manifestations of VZV infection in the CNS in humans.

## Author contribution

THM, CSL, and JLC conceived the idea, CSL identified the patients, MCT, AFH, and SF performed experiments, MCT, AFH, MM, FR, SYZ, THM, SRP, and MC analyzed data, THM wrote the first version of the manuscript, all authors read and approved the final version of the manuscript.

## Acknowledgments

We wish to thank all patients involved in this study. THM is funded by The Independent Research Fund Denmark (4004-00047B) and Aarhus University Research Fund (AUFF-E-2015-FLS-66). AFH received funding from the Faculty of Health, Aarhus University. The Ph.D. scholarship to MCT is funded by the European Union under the Horizon 2020 research and innovation program (H2020) and Marie Skłodowska-Curie Actions–Innovative Training Networks Programme MSCA-ITN GA 675278 EDGE (Training Network providing cutting-EDGE knowlEDGE on Herpes Virology and Immunology). SF is funded by INSERM.

## Declarations

The authors declare that they have no conflict of interest

## References

(1) Gilden D, Cohrs RJ, Mahalingam R, Nagel MA. Varicella zoster virus vasculopathies: diverse clinical manifestations, laboratory features, pathogenesis, and treatment. Lancet Neurol 2009; 8(8):8–731.

(2) Gilden DH, Mahalingam R, Cohrs RJ, Kleinschmidt-DeMasters BK, Forghani B. The protean manifestations of varicella-zoster virus vasculopathy. J Neurovirol 2002; 8 Suppl 2: 75–9.

(3) Gershon AA, Breuer J, Cohen JI, Cohrs RJ, Gershon MD, Gilden D et al. Varicella zoster virus infection. Nat Rev Dis Primers 2015; 1: 15016.

(4) Kircher M, Witten DM, Jain P, O’Roak BJ, Cooper GM, Shendure J. A general framework for estimating the relative pathogenicity of human genetic variants. Nat Genet 2014; 46(3):3–310.

(5) Itan Y, Shang L, Boisson B, Ciancanelli MJ, Markle JG, Martinez-Barricarte R et al. The mutation significance cutoff: gene-level thresholds for variant predictions. Nat Methods 2016; 13(2):2–109.

(6) Duncan CJ, Hambleton S. Varicella zoster virus immunity: A primer. J Infect 2015; 71 Suppl 1:S47–S53.

(7) Ogunjimi B, Zhang SY, Sorensen KB, Skipper KA, Carter-Timofte M, Kerner G et al. Inborn errors in RNA polymerase III underlie severe varicella zoster virus infections. J Clin Invest 2017; 127: 3543–56.

(8) Ablasser A, Bauernfeind F, Hartmann G, Latz E, Fitzgerald KA, Hornung V. RIG-I-dependent sensing of poly(dA:dT) through the induction of an RNA polymerase III-transcribed RNA intermediate. Nat Immunol 2009; 10: 1065–72.

(9) Chiu YH, Macmillan JB, Chen ZJ. RNA Polymerase III Detects Cytosolic DNA and Induces Type I Interferons through the RIG-I Pathway. Cell 2009; 138: 576–91.

(10) Sironi M, Peri AM, Cagliani R, Forni D, Riva S, Biasin M et al. TLR3 Mutations in Adult Patients With Herpes Simplex Virus and Varicella-Zoster Virus Encephalitis. J Infect Dis 2017; 215(9):9–1430.

(11) Bernard G, Chouery E, Putorti ML, Tetreault M, Takanohashi A, Carosso G et al. Mutations of POLR3A encoding a catalytic subunit of RNA polymerase Pol III cause a recessive hypomyelinating leukodystrophy. Am J Hum Genet 2011; 89(3):3–415.

(12) Tetreault M, Choquet K, Orcesi S, Tonduti D, Balottin U, Teichmann M et al. Recessive mutations in POLR3B, encoding the second largest subunit of Pol III, cause a rare hypomyelinating leukodystrophy. Am J Hum Genet 2011; 89(5):5–652.

(13) Nagel MA, Forghani B, Mahalingam R, Wellish MC, Cohrs RJ, Russman AN et al. The value of detecting anti-VZV IgG antibody in CSF to diagnose VZV vasculopathy. Neurology 2007; 68(13):13–1069.

